# Entropy predicts early MEG, EEG and fMRI responses to natural images

**DOI:** 10.1101/2023.06.21.545883

**Authors:** I. Muukkonen, V.R. Salmela

**Affiliations:** Department of Psychology and Logopedics, University of Helsinki, Finland

**Keywords:** visual objects, visual images, image statistics, image entropy, fMRI, EEG, MEG, RSA, DNN

## Abstract

To reduce the redundancy in the input, the human visual system employs efficient coding. Therefore, images with varying entropy (amount of information) should elicit distinct brain responses. Here, we show that a simple entropy model outperforms all current models, including many deep neural networks, in predicting early MEG/EEG and fMRI responses to visual objects. This suggests that the neural populations in the early visual cortex adapt to the information in natural images.

## Main

The principles of information theory have long been applied to the study of visual perception^1^. One influential concept has been the reduction of redundant information, originally suggested by Barlow^2^. It is currently understood that the visual system employs sparse or efficient coding^3,4^ to diminish the redundancy found in natural images^4^. Entropy (*H*), a fundamental concept in information theory^5^, serves as a measure of information that can be obtained from any distribution, such as the luminance histogram (*p*) of an image (1). Entropy is directly associated with redundancy, and lower entropy indicates a higher degree of redundant information. Thus, if the visual system employs efficient coding and economically compresses visual information, images with varying entropy are expected to elicit different patterns of activation. Surprisingly, to our knowledge, this prediction has not been previously tested using neuroimaging data.

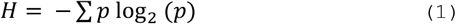

Myriad studies during the last decades have sought to uncover the neural representations of visual objects and the computational models that can explain neural responses to visual stimuli. In a typical experiment, participants are presented with a wide variety of images depicting different objects, and representational similarity analysis (RSA)^6^ is used to compare the neural representations with various models. In RSA, both the data and different models are abstracted into a representational dissimilarity matrix (RDM) containing values of (dis)similarity between all possible stimulus pairs, calculated for example by pair-wise decoding or correlation. The aim is to compare the responses within the representational space and identify the models that best explain the responses across different brain areas (fMRI) or at different time points (M/EEG). The early neural representations, occurring around 100ms (M/EEG) and in the early visual cortex (fMRI), are best explained by models based on image statistics or the early layers of deep neural networks (DNN)^7-13^. Lately, the DNN-based models have generated most enthusiasm. However, image-based models allow easier interpretation of neural representations and serve as a necessary control against which more complex models should be compared.

Here, we compared a simple model derived from image entropy to multiple commonly used models using several independent and openly available datasets that included EEG, MEG, or fMRI data^14-16^. In all of these datasets, participants were presented with several (ranging from 80 to 200) images depicting different objects, such as vehicles (Fig. 1a), animals (Fig. 1b, c), tools, and faces. The images had varying luminance histograms, resulting in entropy values ranging from 5 to 8 bits. Representational dissimilarity matrices (RDMs) were calculated from the data, separately for each subject and timepoint (M/EEG) or region-of-interest (fMRI), and these were then compared to RDMs derived from different model predictions.

First, we compared different image-based models using M/EEG data with visual objects in naturalistic backgrounds (Fig. 1). The image entropy model surpassed all the other image-based models in explaining early M/EEG responses. Significantly higher correlation was found for image entropy than other models and the correlation peaked around 120ms, similarly in MEG (Fig. 1d) and the two EEG (Fig1. e, f) datasets. In addition to image-based models, previous studies have used early layers from DNNs and shown them to match early neural representations reasonably well^8,11,13^. Thus, we next compared the entropy model to predictions derived from pre-trained DNNs, specifically different layers from AlexNet^17^, ResNet50^18^, and CLIP-ResNet50^19^. As expected, the DNN-models showed relatively high correlations with the neural data, surpassing all commonly used image-based models. Remarkably, the simple and parameter-free entropy model was comparable with the complex DNN-models. In the MEG (Fig. 1g) dataset and the other EEG (Fig. 1i) dataset, entropy and the best-performing DNN correlated equally well with the data. In the other EEG dataset, entropy showed significantly higher correlation than the best DNN (Fig. 1h).

**Figure 1.**
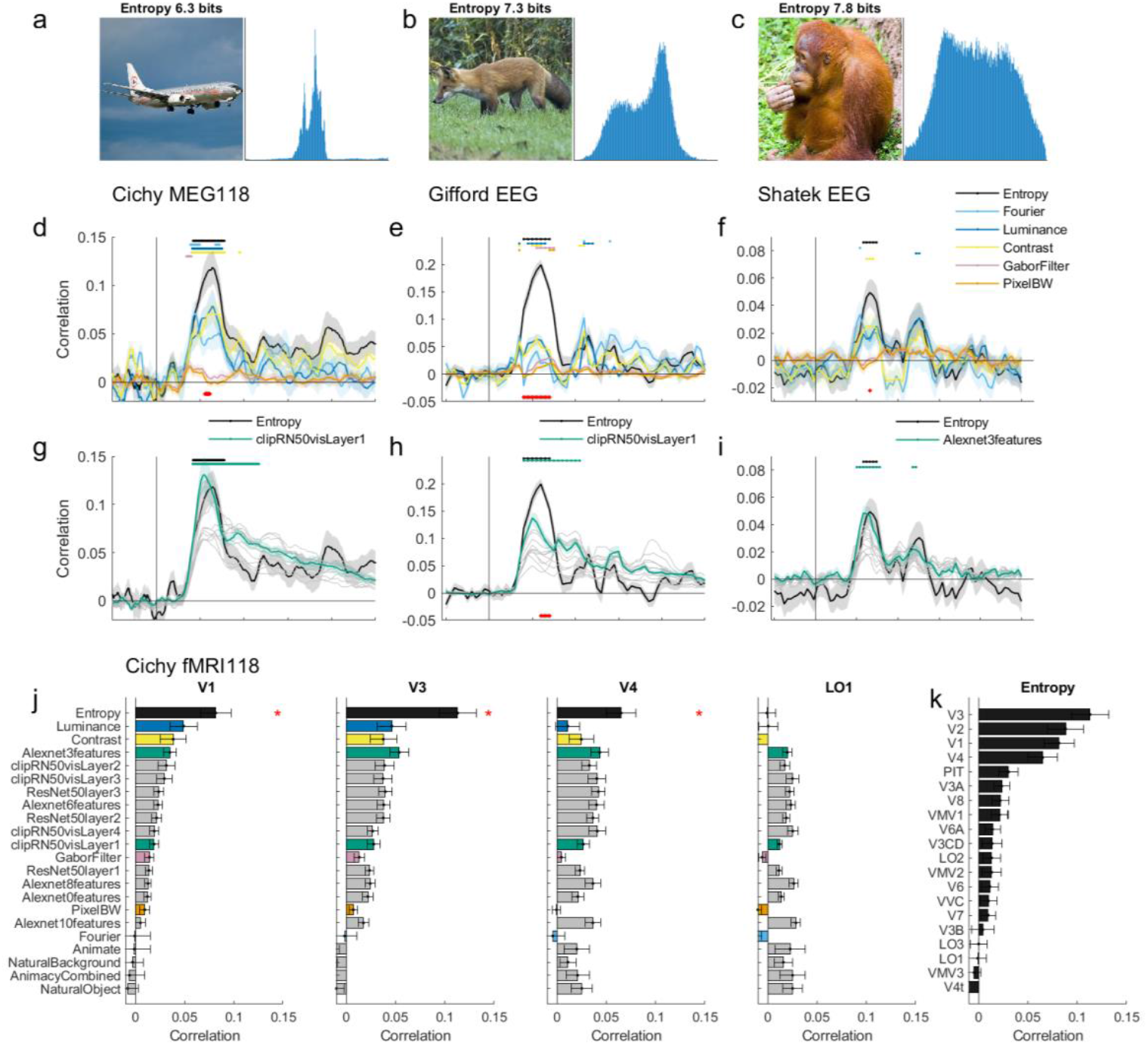
Examples of stimulus images and correlations between model-RDMs and data-RDMs. **a-c**) Three example images with different luminance histograms and entropies (from Cichy118). The histogram has 256 bins, and therefore maximum entropy is 8 bits. **d-f**) Correlation of entropy vs. other image-based models in three datasets. Red dots on the bottom of the plot indicate timepoints when entropy significantly outperformed other models (permutation test, p<0.05). Colored dots on the top of the plot indicate timepoints where each model correlation was significantly above zero (permutation test, p<0.05). **g-i**) Entropy compared with DNNs. The best DNN is shown in green, other DNNs are shown in gray. **j**) Model correlations in fMRI data in areas V1, V3, V4 and LO1 (HCP parcellation). The data is sorted according to V1. The red asterisk indicates areas where entropy was significantly better than all other models (permutation test, p<0.05). **k**) Correlations of the entropy model in different visual areas. In d-e, shaded areas show SEMs. In j-k error bars depict SEMs.

In fMRI, the results were similar. In the early visual areas V1-V4, the entropy model had a significantly higher correlation with the data than any other model (Fig. 1j). The correlation of entropy and the neural data gradually decreased along the visual processing hierarchy (Fig. 1k). Neural representations in higher visual areas, such as the lateral occipital cortex (LO1), did not correlate with entropy and were instead best explained by DNNs and the naturalness (e.g., animal vs. tool) of the objects. Thus, representations related to entropy in the brain seem to be highly confined to the early visual cortex and to a relatively short time window (80 – 160ms). This is in contrast with the predictions from DNNs, which correlate with both early and higher areas, and across a longer time window. Thus, image entropy allows for easier interpretation of the early neural representations by being both a simpler model as well as having more spatiotemporally localized neural correlates.

One notable distinction between image entropy and most of the other models is that entropy relies on global image statistics, that is, it is insensitive to the spatial arrangement of the pixels. Consequently, one explanation for the success of entropy is that it uses global image statistics, whereas models relying on local information (e.g., pixel, Gabor filter) fail when natural images are used as stimuli. Other models based on information from the whole image, root-mean-square (RMS) contrast, mean luminance, and global Fourier spectrum, were better than local image-based models but still worse than entropy (Fig. 1d-f). Models based on the kurtosis and skewness of the luminance histogram were equal or worse than entropy. In fMRI studies, comparing (Fourier) phase scrambled and intact objects is a common way to functionally localize higher-level visual areas such as FFA^20^. This is in accordance with entropy, as pixel-wise scrambled naturalistic images should elicit similar responses as intact images in early visual areas.

Notably, results differed substantially in one of the analysed datasets (Cichy92, both MEG and fMRI). In this dataset, the stimuli contained no backgrounds (Fig. 2a), meddling with the natural image statistics and introducing artificial object boundaries. Interestingly, the local rather than the global models showed higher correlations with the data (Fig. 2b). The global entropy model performed worse than the local models, and the silhouette model best explained the early responses. Similarly, in fMRI, the neural responses to stimuli without backgrounds were best explained by local models (Fig 2d). Although entropy performed worse than other models, it still showed relatively similar results as in the other datasets: it correlated more with early than higher visual areas in fMRI (Fig. 2e) and peaked at around 100ms in MEG (Fig. 2b, c).

**Figure 2.**
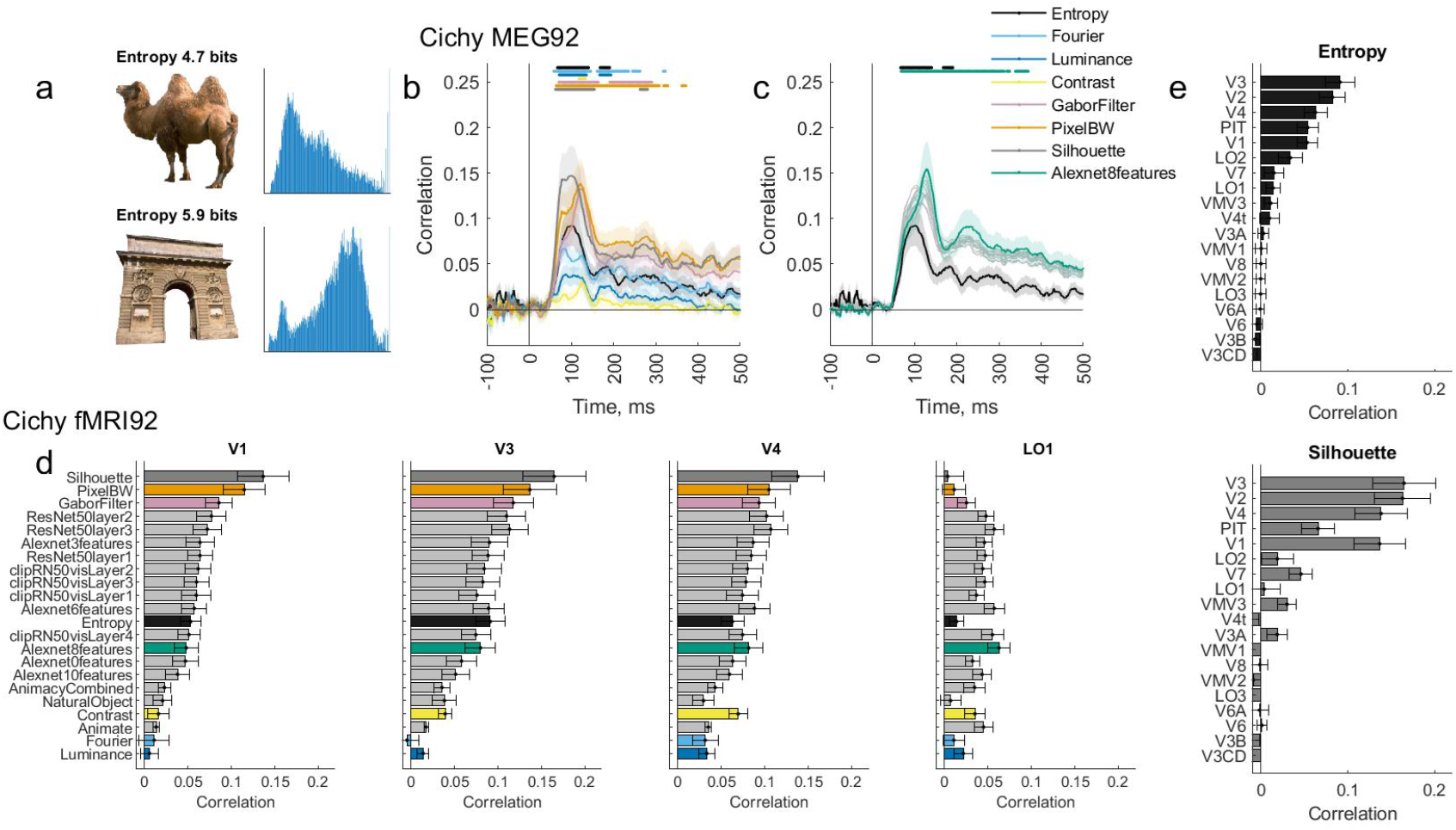
Entropy with images without background. **a**) Two example images and their histograms. The histograms have a high peak of white pixels that is cut off. **b-c)** Entropy vs. other models in the MEG dataset. The best DNN is shown in green, other DNNs in gray. The dots above the images represent timepoints with significantly higher than zero correlation (permutation test, p<0.05), and the shaded areas indicate SEMs. **d**) Entropy vs. other models in fMRI areas V1, V3, V4 and LO1 (HCP parcellation). The data is sorted according to V1. **e**) Entropy and silhouette model correlations in different visual areas. In d-e error bars indicate SEMs.

Finally, we compared the models with each other. The entropy model correlated only weakly with the other models in all four datasets (Fig. 3a-d), showing the highest correlations with contrast (0.25 – 0.35) in datasets with backgrounds (Fig. 3a-c), and somewhat higher correlations in the Cichy92 dataset without backgrounds (Fig. 3d; luminance: 0.47; clipRN50 layers 2-4: 0.5 – 0.52). This suggests that the entropy model captures a previously unexplained portion of the M/EEG and fMRI representations. Accordingly, Figures 3e-h show the independent contribution of each model in explaining the variance of the data. Specifically, including the entropy model increases the variance explained compared to a regression model containing all the other models except for entropy. This was also true to a lesser extent for the dataset without backgrounds (Fig. 3h, Cichy92). Including an entropy model, therefore, allows for a better explanation of neural data, even when DNNs are also used.

**Figure 3.**
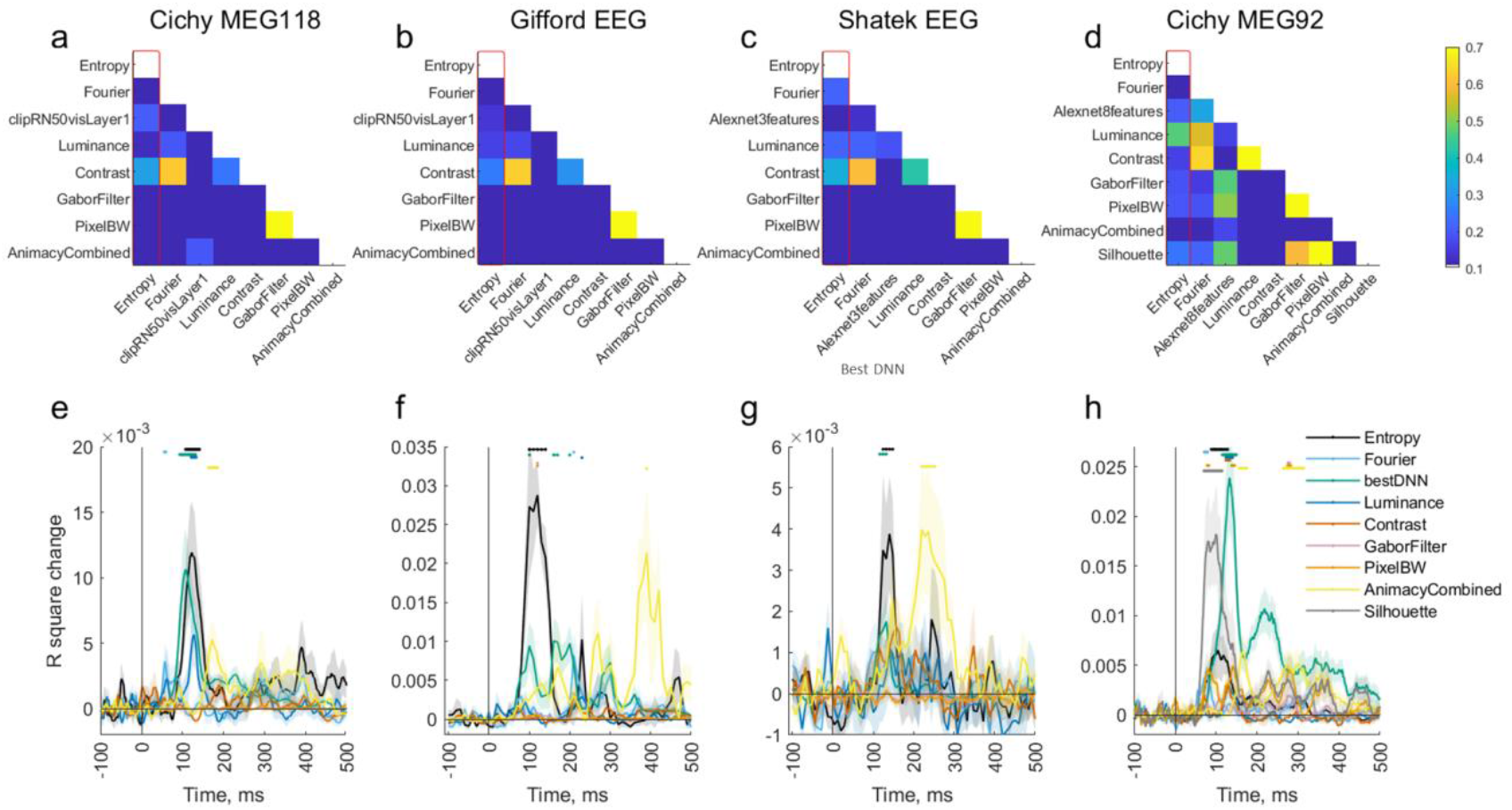
Correlations between models and the results of the regression analysis. **a-d**) Correlations between models in the four datasets. Red rectangle highlights the correlations of entropy with other models. **e-h**) Change in R2 value in leave-one-model-out comparison in the four datasets. The change in R2 value indicates independent contribution of each model in explaining the variance in the data. Dots above images show significant timepoints (permutation test, p<0.05), shaded areas depict SEMs.

In this study, we demonstrated that image entropy best explains early M/EEG and fMRI response to visual images in several independent datasets. To our knowledge, the role of image entropy in explaining brain responses to visual objects has not been previously tested. The entropy model explained well both the early M/EEG and early visual cortex fMRI responses to visual objects in naturalistic backgrounds. It outperformed conventional image-based models and showed comparable or superior performance compared to basic DNN models. The role of entropy and other global models was specific to stimuli with naturalistic backgrounds, while local models explained the responses to stimuli without backgrounds better. Image entropy correlated only weakly with the other models and consequently explained a part of the neural representations not captured by previous models.

Information theory has been applied to behavioral and physiological vision research for decades^1-4^, and it has been proposed as a better framework for the interpretation of the results from multivariate neuroimaging experiments^21,22^. For example, the classification accuracy from multivariate decoding analyses is pointed out to be an estimate of information^21^. Here we provide evidence for this notion and show the early neural responses to be highly sensitive to the variation of entropy between stimuli. Our results thus suggest that the redundancy in visual images is efficiently reduced in the visual system, and the resulting representations are measurable with MEG, EEG and fMRI. This adjustment to the amount of information in the visual stimuli can either be inherited from pre-cortical processes or achieved in the early visual areas in order to attain stable and independent population responses^23^. The results also imply that including image entropy as a predictor of neural responses can increase the share of neural representations we are able to explain, serve as a better control or benchmark than commonly used image statistics, and, due to its simplicity, offer more tractable explanations of neural computations.

## Methods

### Datasets

We used four openly available datasets, two using both MEG and fMRI, and two using EEG, and all having images of objects as stimuli. We refer the reader to the original articles for the specific details of data collection and preprocessing. Hereafter, we refer to the datasets as *Cichy118*^14^ (experiment 2), *GiffordEEG*^15^ (test set), *ShatekEEG*^16^, and *Cichy92*^14^ (experiment 1, session 1). The number of subjects was 15, 10, 24, and 16, respectively; number of stimuli types 118, 200, 80, 92; M/EEG channels 306, 17, 24, 306; sampling rate 1000Hz, 100Hz, 250Hz, 1000Hz; trials per stimulus type 20 – 30, 80, 16, 10 – 15; stimuli were shown for 500ms, 100ms, 100ms, 500ms; and stimulus onset asynchrony was 0.9 – 1s, 200ms, 50ms, 1.5 – 2s. Both EEG datasets used a rapid serial visual presentation paradigm design (RSVP)^24^. From *ShatekEEG*, we used only the occipital and parietal-occipital channels and one-fifth of the total 400 stimuli, 2 from every category (image numbers 1 and 10).

## Analysis

All data representational dissimilarity matrices (RDM)^6^ were created separately for each subject and each timepoint in M/EEG or region of interest (ROI) in fMRI. As a dissimilarity metric, either one-minus-correlation (*GiffordEEG*, fMRI-data) or support vector machine (SVM) decoding accuracy (*CichyMEG118, CichyMEG92, ShatekEEG*) were used. For *CichyMEG118* and *CichyMEG92*, SVM-accuracies were provided by the original authors, and for *ShatekEEG* they were calculated with DDTBOX^25^ which applies LIBSVM^26^, with 16ms sliding window, 8ms step-size, 2-fold cross-validation (trials randomized to folds) repeated 4 times, and normalizing data before decoding. The SVM used the M/EEG channels as features, and in correlation, vectors of channel or voxel values were correlated between all possible stimulus pairs after averaging trials for each stimulus type. In fMRI, ROIs were defined with the HCP-parcellation^27^.

The image-based model RDMs were created as follows. Color images were first transformed to grayscale and resized to 256*256 pixels. Image entropy was calculated by Matlab’s *entropy()* –function, which takes the histogram *p* of pixel values (with 256 bins), removes zero values, normalizes the values of *p* to have a sum of 1, and calculates entropy (-sum(p.*log2(p))). Fourier model uses the two-dimensional Fourier transform of the image matrix (*fft2* -function in Matlab). Pixel model is based on the vectorised raw values of all pixels of the images. Gabor filter model is based on a bank of Gabor filters. Each image was filtered with 36 filters (6 different spatial scales at 6 different orientations). The center spatial frequency of the filter was varied from 1 to 64 cycles/image width. The spatial frequency bandwidth of the filters was one octave and the orientation bandwidth was 30°. Luminance is the mean of all pixel values, and contrast is root mean square contrast (RMS), namely the standard deviation of pixel values divided by the mean of all pixels. For all models, the RDM was then calculated by taking either the 1-correlation of the vector of values or, in entropy, contrast and luminance models, the absolute difference between the values representing each stimulus, corresponding to one-dimensional Euclidean distance.

DNN models were created with the Net2Brain Toolbox^28^, which contains several pretrained DNN models and inbuilt functions to derive RDMs to any set of images from the DNNs. We used three pretrained DNNs, AlexNet^17^, ResNet50^18^, and CLIP-ResNet50^19^. These were selected as the first two are used as the examples in the toolbox’s manual and are widely used, and the third was shown to be best predictor of fMRI-data in V1^27^. We utilized the default layers provided by the toolbox: Alexnet: 0, 3, 6, 8 and 10 features; ResNet50: layers 1, 2, and 3; clipResNet50: visLayers 1, 2, 3, and 4.

After creating the RDMs, they were vectorised (lower triangle) and correlations (Spearman) between models and data were calculated separately for each timepoint/ROI and subject. To test the individual contribution of different models, we conducted regression analyses with all models as predictors as well as leaving one model out, and compared whether the R^2^ of the regression model increased when a given model was included. Means across subjects are plotted, and statistical significance was determined with permutation tests^29^. The signs of the model correlation values (positive or negative) for each subject were randomly shuffled 1000 times, over-subjects mean model-data correlation was calculated for each permutation fold, and the highest correlation value (from all timepoints) was taken. This generates a permutation distribution, adjusted for multiple comparisons across timepoints. This distribution was compared to the true mean correlations, which were defined significant if they were higher than 95% of the shuffled values (p<.05, one-sided). In the regression analyses, the same procedure was used after first correcting all the values (separately for each subject and model) with the mean of the R^2^-increase in the baseline period (−100 – 0ms), providing an empirical estimate of the R^2^-increase during a period of (presumably) pure noise in the data.

## Data and code availability

All data are publicly available at: *Cichy92 & Cichy118*: http://userpage.fu-berlin.de/rmcichy/fusion_project_page/main.html; *GiffordEEG*: https://doi.org/10.17605/OSF.IO/3JK45; *ShatekEEG*: https://openneuro.org/datasets/ds003887. A summary of data and scripts to conduct the analyses and reproduce the results and figures of the current study are available at https://osf.io/q2cze/.

## Notes

### Competing Interest Statement

The authors have declared no competing interest.

### Summary of Updates

link to code/data provided referencing style changed some minor edits

